# Species rich but data poor: leveraging distribution modelling techniques to map cetacean occurrence in South Asian waters

**DOI:** 10.1101/2024.12.27.630491

**Authors:** Imran Samad, Dipani Sutaria, Kartik Shanker

**Author notes:** Corresponding author: KS Lab, CES, IISc., Bangalore, India – 560012.

## Abstract

Cetaceans play important ecosystem roles but are challenging to study since they are highly mobile and spend their entire life at sea. More than a third of all cetaceans occur in South Asian waters where they overlap with several key threats including fisheries, pollution, and high vessel traffic. However, baseline information on their occurrence at the sub-continental scale is lacking even for relatively well studied species. To address this gap, we collated sighting information on cetaceans from the North Indian Ocean region (5-25°N, 65-95°E) and used species-tuned distribution models and ensemble modelling approaches with physiographic and oceanographic predictors to map habitat suitability and species richness. We used Multivariate Environmental Similarity Surfaces (MESS) indices and expert reviews to validate our predictions and better infer species distribution. We collated 2329 sighting records for 31 species and modelled the distribution of 18 species. Areas of high species richness, defined by high slope and bathymetric complexity, occurred along the east coast of India, in south Sri Lanka, and around the Lakshadweep and Andaman and Nicobar archipelagos. Species richness patterns were generally consistent with the six identified ecoregions in the area. Most of our predictions were given high scores by the expert reviewers. Our maps underpredicted the occurrence of a few oceanic species, highlighting the need for focused offshore surveys, and overpredicted areas with high MESS indices for all species. We discuss the occurrence patterns and their drivers for these cetacean species while highlighting knowledge gaps and the importance of fine-scale distribution data for conservation planning, especially during the development of regional management plans for cetaceans.

## Introduction

Owing to their diversity, top trophic positions, large-scale distributions, and long life-histories, cetaceans play crucial roles in ecosystem function like nutrient transport and top-down control (Chami et al., 2019; Kiszka et al., 2022a; Gilbert et al., 2023). Their occurrence, movement, and distribution may reflect complex ecosystem processes and changes in biogeochemical oceanography and prey distributions due to climate change as they can migrate from unfavourable habitats (Henson et al., 2017). Consequently, cetaceans can serve as sentinels of change (Moore 2008; Bossart 2010; Hazen et al., 2019). For example, southern right whales (*E. australis*) have been increasingly observed using higher latitudes for foraging over the past few decades likely due to changes in ocean fronts and productivity (Derville et al., 2023). Moreover, changes in cetacean distribution and abundance can also have cascading effects on the environment, emphasizing the need to better understand their ecology. However, their offshore and wide-ranging nature also impedes our understanding of their distribution, especially in the high seas and in developing countries where most species occur (Kaschner et al., 2011; Kaschner et al., 2012).

In the past century, human impacts on the sea have grown exponentially and more than 25% of all cetacean species today are threatened with extinction (Braulik et al., 2023). Primary threats to cetaceans include anthropogenic activities like fisheries, pollution, and shipping (Reeves et al., 2005; Avila, Kaschner, & Dormann, 2018; Kiszka et al., 2022b), with their impacts prevalent in at least half of all critical cetacean habitat globally (Avila, Kaschner, & Dormann, 2018). Regions in South and South-East Asia are identified as high-risk habitats (Braulik et al., 2023), but also support more than half of all cetaceans globally (Pompa, Ehrlich, & Ceballos, 2011). Yet very little systematic information exists about the broad-scale ranges of different species occurring in this region (Jefferson et al., 2014; Sahri et al., 2021), particularly from the Northen Indian Ocean in the coastal and offshore waters of South Asian countries. These include the territorial and Exclusive Economic Zones of at least five countries i.e., India, Pakistan, Sri Lanka, Bangladesh, and Myanmar, covering an area of more than 3.5 million km^2^. Here, human impacts are some of highest globally (Halpern et al., 2008; Hou et al., 2015; Meijer et al., 2021; Welch et al., 2022) and with a potential impact on several cetacean species (Anderson et al., 2020; Elliot et al., 2024).

More than 25 species of cetaceans have been recorded from South Asian waters mainly based on stranding reports of which a few like humpback dolphins (*Sousa sp.*), Indo-pacific finless porpoises (*N. phocaenoides*), are resident in nearshore coastal waters all along the Indian coast, while species such as bottlenose dolphins (*Tursiops sp.*), spinner dolphins (*S. longirostris*) and spotted dolphins (*S. attenuata*) also occur close to the coastline along the east coast given the deeper waters close to shore in the east (Kumarran 2012, MMRCNI 2024).

Interview-based studies (Sule et al., 2017; Karnad 2022) and stranding records (Kumaran 2002; Ilangakoon 2006; Dudhat et al., 2022) have highlighted the presence of several cetacean species, but boat-based survey efforts to understand their distribution have been sporadic and opportunistic (Singh 2003; Sutaria & Jefferson, 2005; Sule et al., 2016; Panicker 2020). Even for relatively well studied species like humpback dolphins and the Indo-pacific finless porpoise (Sutaria et al., 2015; Sule et al., 2017), a systematic assessment of their distribution is lacking, except near Pakistan and northwest India (Kiani et al., 2023). Reports on larger species like humpback whales (*M. novaeangliea*) (Anderson et al., 2022), blue whales (*B. musculus*) and other baleen whales (*Balaenoptera sp.*) (Sutaria et al., 2016; Sutaria et al., 2017) have improved our understanding of their migration patterns while providing insights into the threats faced by different species. Anthropogenic activities like fisheries interactions (Yousuf et al., 2009; Joseph et al., 2021), gear entanglement and bycatch (Anderson et al., 2020), and noise and pollution (Kumarran 2012) are primary threats faced by cetaceans in the region. While the Indian government has created an action plan to conserve dolphins in the region (GoI, 2021), a lack of basic knowledge of their distribution hinders any conservation effort and excludes cetaceans when management measures and Environmental Impact Assessments (EIAs) are being carried out for proposed development plans.

Species Distribution Models (SDMs) or Environmental Niche Models have gained popularity in understanding and mapping the distribution of species based on their associations with environmental parameters (Elith and Leathwick 2009). Their ability to work with a limited, non-systematic, and presence-only data at multiple scales to provide accurate information on species-habitat associations and distributions (Fois et al., 2018) makes them a robust choice for understanding rare and data deficient species, including those in marine ecosystems (Robinson et al., 2017; Melo-Merino et al., 2020). A suite of algorithms and modelling choices (Li & Wang 2012; Valavi et al., 2022) along with pragmatic guidelines and suggestions (Robinson et al., 2017; Soley-Guardia et al., 2024) provides users with the flexibility to fine-tune modelling approaches to the idiosyncrasies of their own data and mitigate challenges associated with low sample sizes and geographic bias in presence records (Grimmett et al., 2020; Radomski et al., 2022)..Consequently, SDMs have found application in identifying conservation priority and survey areas (Rana et al., 2022), predicting range shifts under climate change scenarios (Austin & Niel 2010), mapping species diversity (Sahri et al., 2021), informing EIAs and developing management plans (Sofaer et al., 2019)

In this study, we use SDMs to map the habitat suitability of 18 marine cetaceans occurring in the Arabian Sea and the Bay of Bengal. In addition to a suite of modelling approaches, we solicited expert reviews to verify and validate the maps. We then identified areas where different species may occur to estimate and demarcate areas of high species richness in the region. Finally, we use our results to discuss gaps in our knowledge of species occurrence, areas to focus survey efforts on, and potential strategies for conserving marine cetaceans in South Asia.

## 2. Methods

### 2.1. Study area

We predict the distribution of cetaceans in the northern Indian Ocean between 5-25°N and 65- 95°E. The Indian subcontinent divides the northern Indian Ocean basin into the Arabian sea to the west and the Bay of Bengal to the east. This area can be divided into six ecoregions (four along the mainland and two archipelagos) based on topographic features and species biogeography (Spalding et al., 2007).

On the northwest coast of the Indian subcontinent, the continental shelf is wide and extends 150-200 km into the Arabian sea compared to the southwest coast where it is much narrower, extending to only half the distance. On the east coast the shelf extends between 20 km and 100 km and has a steeper slope. Depth and contour profile also varies on both continental shelves and ranges up to 300 meters.

The Arabian Sea and the Bay of Bengal are also characterised by seasonal upwellings and high productivity, especially in the coastal regions (Panikkar & Jayaraman 1966). The Andaman and Nicobar islands lie at the southeast end of the study area between the Bay of Bengal and the Andaman sea while the Lakshadweep islands lie around the same latitude but much closer to the Indian mainland in the southeast Arabian sea. Both the archipelagos are characterised by high depth and complex seafloor geomorphology. Around the island nation of Sri Lanka, the continental shelf is much narrower, extending only 15-30 km into the sea before the onset of the continental slope.

### 2.2. Collating and processing species sighting data

We collected data on sighting geo-coordinates of cetaceans in South Asian waters from multiple sources. Firstly, we searched for published reports and peer-reviewed papers using keywords including ‘cetaceans’, ‘dolphin’, ‘whale’, and ‘India’, ‘Bangladesh’, ‘Pakistan’, ‘Sri Lanka’ between 2000 and 2024 and screened for documents where GPS coordinates of species sighting were provided. If sightings were marked on maps, they were geo-referenced based on latitudinal and longitudinal grids on the map to extract coordinates. Secondly, we searched for sighting records on public databases like the Marine Mammals of India website (MMRCNI 2024) and GBIF. Thirdly, we accessed sighting information from social media platforms and local researcher networks. After compiling all records, we identified and removed a) duplicates based on the similarity of species sighted, location, and time, and b) records with uncertain or inaccurate species identity.

Opportunistically collected data is likely to be spatially biased due to uneven sampling effort which should be explicitly accounted for when modelling species distributions (Phillips et al., 2009; Jeliazkov et al., 2022; Baker et al., 2024). To minimize the impact of sampling bias, we filtered out sighting locations in close proximity (Kramer-Schadat et al., 2013; Inman et al., 2021) for all species at two different scales i.e., 10km and 100km, using the *spThin* package (Aiello-Lamens et al., 2015) in R (Table 1). Only species with at least 10 sighting records, distributed across a wide range of their known occurrence regions were used for analysis (Stockwell & Peterson, 2002).

**Table 1:** A summary of species sighting records in the study region. Data for the two species of humpback dolphins, and the two species of bottlenose dolphins were combined and modelled as two species. ‘Sightings’ refer to the raw number of data points and ‘Filtered sightings’ refer to the number of data points left after filtering (or spatial thinning). Optimal ‘Filtering scale’ (10 vs 100) was obtained for each species based on model performance. Classification was based on the known ecology and occurrence of each species in the area, and few oceanic species also tend to occur close to the coast.

| Species | Scientific name | IUCN status | Classification | Sightings | Filtered sightings | Filtering scale (km) |
| --- | --- | --- | --- | --- | --- | --- |
| Indo-Pacific Humpback dolphin | *Sousa chinensis* | Vulnerable | Coastal | 470 | 157 | 10 |
| Indian Ocean Humpback dolphin | *Sousa plumbea* | Endangered | Coastal | 155 |  |  |
| Blue whale | *Balaenoptera musculus* | Endangered | Oceanic | 437 | 14 | 100 |
| Spinner dolphin | *Stenella longirostris* | Least Concern | Oceanic | 374 | 171 | 10 |
| Indo-Pacific Bottlenose dolphin | *Tursiops aduncus* | Near Threatened | Oceanic | 203 | 58 | 100 |
| Common Bottlenose dolphin | *Tursiops truncatus* | Vulnerable | Oceanic | 37 |  |  |
| Sperm whale | *Physeter macrocephalus* | Vulnerable | Oceanic | 158 | 33 | 100 |
| Humpback whale | *Megaptera novaeangliae* | Least Concern | Oceanic | 63 | 12 | 100 |
| Bryde's whale | *Balaenoptera edeni* | Least Concern | Oceanic | 59 | 15 | 100 |
| Irrawaddy dolphin | *Orcaella brevirostris* | Critically Endangered | Coastal | 59 | 7 | 100 |
| Striped dolphin | *Stenella coeruleoalba* | Least Concern | Oceanic | 58 | 36 | 100 |
| Long-beaked Common dolphin | *Delphinus delphis* | Least Concern | Oceanic | 54 | 22 | 100 |
| Short-finned Pilot whale | *Globicephala macrorhynchus* | Least Concern | Oceanic | 39 | 30 | 10 |
| Pan-tropical spotted dolphin | *Stenella attenuata* | Least Concern | Oceanic | 26 | 23 | 10 |
| Risso's dolphin | *Grampus griseus* | Least Concern | Oceanic | 23 | 18 | 10 |
| Killer whale | *Orcinus orca* | Data Deficient | Oceanic | 21 | 9 | 100 |
| False Killer whale | *Pseudorca crassidens* | Near Threatened | Oceanic | 17 | 13 | 10 |
| Indo-Pacific finless porpoise | *Neophocaena phocaenoides* | Vulnerable | Coastal | 17 | 11 | 100 |
| Dwarf Sperm whale | *Kogia sima* | Least Concern |  | 6 |  |  |
| Fraser's dolphin | *Lagenodelphis hosei* | Least Concern |  | 5 |  |  |
| Melon-headed whale | *Peponocephala electra* | Least Concern |  | 5 |  |  |
| Rough-toothed dolphin | *Steno bredanensis* | Least Concern |  | 5 |  |  |
| Fin whale | *Balaenoptera physalus* | Vulnerable |  | 4 |  |  |
| Pygmy Sperm whale | *Kogia breviceps* | Least Concern |  | 3 |  |  |
| Longman's Beaked whale | *Indopacetus pacificus* | Least Concern |  | 2 |  |  |
| Minke whale | *Balaenoptera acutorostata* | Near Threatened |  | 2 |  |  |
| Omura's whale | *Balaenoptera omurai* | Data Deficient |  | 2 |  |  |
| Pygmy Killer whale | *Feresa attenuata* | Least Concern |  | 2 |  |  |
| Blainville's Beaked whale | *Mesopledon densirostris* | Least Concern |  | 1 |  |  |
| Cuvier's Beaked whale | *Ziphus cavirostris* | Least Concern |  | 1 |  |  |
| Pygmy Blue whale | *Balaenoptera musculus brevicauda* | Data Deficient |  | 1 |  |  |
| Baleen whale |  |  |  | 14 |  |  |
| Small delphinid |  |  |  | 5 |  |  |

When only presence data is available, species distribution models require background data to be extracted from the study area, against which the properties of locations with sighting records are contrasted to understand the habitat suitability of species (VanDerWal et al., 2009; Breen et al., 2016). The approach and extent used to sample background data can greatly influence model outcomes (VanDerWal et al., 2009; Rana et al., 2024; Whitford et al., 2024) and therefore must be guided by species ecology and the region of interest (Saupe et al., 2012). One such approach that minimizes the impact of sampling bias (Inman et al., 2021) and increases model sensitivity (Barbet-Massin et al., 2012) is sampling background points only from regions within a specified distance around presence locations. This buffer distance should be such that it reflects areas that are accessible to the species through dispersal (Merow et al., 2013). Based on preliminary analysis, we chose a buffer distance of 1 degree for coastal species i.e., Humpback dolphins, the Irrawaddy dolphin (*O. brevirostris*), and the Indo-pacific finless porpoise and 2 degrees for oceanic species to define the background. For each species, we sampled 10,000 points from the background that were used as pseudo-absence points for further analysis.

### 2.3. Collating predictor variable data

To model cetacean distributions, we selected physiographic and oceanographic variables based on our knowledge of species from previous studies (Stephenson et al., 2020; Sahri et al., 2021). These included depth, slope, distance to mainland, current velocity, primary productivity, mean SST, annual temperature range, and the presence of temperature fronts. To avoid model overfitting, we removed variables that were highly correlated (Pearson’s r > 0.7) and all remaining variables were used to model species distributions. Temperature fronts were only observed >20 km away from the coast and so were not used to model the distribution for coastal species (details on the variables used, their descriptions and sources are listed in Table 2). All rasters were resampled at a resolution of the coarsest raster (1/12 degrees) and their values were extracted for sighting and background locations. Maps of all variables used to model distributions can be found in Figure A1.

**Table 2:** List of variables used to model the distribution of all species. Highly correlated variables (Retained = N) were not used in the analysis.

| Category | Variable | Description | Source | Retained |
| --- | --- | --- | --- | --- |
| Physical | Distance to mainland | Distance of each pixel to the coastline of the Indian subcontinent | Derived | Y |
|  | Distance to river mouth | Distance of each pixel to the closest river mouth | Derived from OSM river map | N |
|  | Depth | Bathymetric depth | GEBCO | Y |
|  | Slope | Bathymetric slope | Derived from the depth layer | Y |
|  | Seabed roughness | Index for abrupt changes in depth | Derived from the depth layer | N |
| Oceanographic | Current velocity | Mean annual speed of ocean current flow in m/s | Bio-Oracle | Y |
|  | var(Current velocity) | Annual variation in current flow | Bio-Oracle | N |
|  | Primary productivity | Mean annual production in g/year | Bio-Oracle | Y |
|  | var(Primary productivity) | Annual variation in production | Bio-Oracle | N |
|  | Mean temperature (SST) | Mean annual sea surface temperature | Bio-Oracle | Y |
|  | Annual temperature range | Range (max-min) in annual temperature | Bio-Oracle | Y |
|  | Temperature fronts | Index for abrupt changes in SST | Derived from the SST layer | Y |

### 2.4. Species distribution modelling

We build species-tuned distribution models to a) predict habitat suitability for each species and selected the top model based on different metrics and b) map species occurrence based on the top habitat suitability model and various thresholding criteria. We then assessed our predictions by comparing them with maps chosen by reviewers, based not on evaluation metrics but their knowledge of the species.

#### 2.4.1. Model tuning and selection

Several techniques have been developed to predict the distribution of rare and data deficient species using presence-only data (Aguirre-Gutiérrez et al., 2013; Radomski et al., 2022; Valavi et al., 2022), and ensembles of multiple models are typically known to provide high predictive performance and stability (Montoya-Jimnez et al., 2022; Valavi et al., 2022), to predict distributions. To build ensemble models for all species, we chose six different algorithms based on regression, tree-based, and machine learning techniques. These included Generalised Additive Models (GAM), Multivariate Adaptive Regression Spline (MARS), balanced Random Forest (RF), Boosted Regression Tree (BRT), Support Vector Machine (SVM) and MaxEnt. Model hyperparameters for all algorithms but GAMs were tuned for each species to obtain optimal predictions (Merow et al., 2014; Hallgren et al., 2019; Valavi et al., 2022) using the ‘caret’ and ‘ENMeval’ (Muscarella et al., 2014) packages.

These optimised parameters were used to build predictive models for each species using a 5- fold spatial cross validation approach to avoid model overfitting (Soley-Guardia et al., 2024). The study area was first partitioned into grids of size 1200X1200 km^2^ based on the spatial autocorrelation of all rasters estimated using the ‘blockCV’ package (Valavi et al., 2018). Five folds were randomly assigned to these grids of which four were used to build a model and the 5^th^ one was used to test it. This process was replicated five times and average values of test AUC scores were estimated to assess model performance. Models with a high difference (>0.1) between training and test AUC values were removed and AUC weighted ensembles of a) all models, and b) top 3 models based on were created. Altogether, for each species, eight habitat suitability maps (six algorithms and two ensembles) were generated and compared. All models were run using the ‘sdm’ package (Naimi & Araujo 2016).

#### 2.4.2 Assessing model performance

While AUC scores have been traditionally used to assess the predictive performance of species distribution models, they may not be sufficient to evaluate models build using presence-only data because of potentially high commission error (Soley-Guardia et al., 2024). An alternative way of assessing the performance of models build using presence only data is to use the Continuous Boyce Index or CBI (Hirzel et al., 2006; Boyce et al., 2002) compares the model to a random or NULL model. We estimated CBI values for all models using the ‘ecospat’ package (Di Cola et al., 2017) and the best model of the eight for each species was selected based on high AUC and CBI values. In case top models had very similar CBI values, the model predicting a more accurate distribution from what is known of the species was chosen.

#### 2.4.3. Predicting species presence and mapping richness

Species presence-absence (PA) maps can be created by using a threshold for habitat suitability predictions which are based on varying criteria. We used three different thresholds that a) maximised the sum of specificity and sensitivity, b) provided equal values of specificity and sensitivity, and c) selected areas where species were more likely to occur than random (Hirzel et al., 2006) to generate three different Presence-Absence (PA) maps from the best habitat suitability model prediction for each species. These maps were visually assessed to select one that most closely described the known and expected distribution of each species. Since humpback whales do not occur in the Bay of Bengal, their distribution was restricted to the Arabian sea and the Gulf of Mannar. PA maps for all species were aggregated to create species richness maps in the study area.

#### 2.4.4. Validating predictions

We assessed uncertainty in habitat suitability model predictions by estimating the Multivariate Environmental Similarity Surfaces or MESS index (Elith et al., 2010) to map areas with environmental conditions beyond the range of that at sampled locations i.e., areas where habitat suitability is extrapolated. We also assessed the accuracy of our models by reaching out to cetacean researchers with in-depth knowledge of the species in the study area to seek their opinion on our predictions. Specifically, reviewers were asked to (a) select the best (of the eight) habitat suitability model for each species based on their understanding of species ecology and the study area, and provide an overall score of the prediction’s accuracy; (b) select the threshold that best describes where species may be found in the region based on the habitat suitability model selected in step (a); and (c) mark areas with false-positive and false-negative predictions in the PA map selected in step (b). Finally, we qualitatively assessed the differences between predictions from our top models and the models chosen by reviewers.

## 3. Results

### 3.1. Model performance

We collated 2329 sighting records of 31 species from the study area. Several records for the two species of bottlenose dolphins (*T. aduncus* and *T. truncatus*) and the two species of humpback dolphins (*S. plumbea* and *S. chinensis*) were uncertain in species identity and so were merged to form two species groups i.e., bottlenose dolphins (*Tursiops sp.*) and humpback dolphins (*Sousa sp.*). Records where species identity could not be established with certainty (n=19) were removed from further analysis. Of the 31 species, only 18 had sufficient data to model their distribution. These included humpback dolphins, bottlenose dolphins, spinner dolphins, striped dolphins, pan-tropical spotted dolphins, long-beaked common dolphins, Irrawaddy dolphins, Risso’s dolphins, short-finned pilot whale, Indo-pacific finless porpoises, killer whales, false killer whales, sperm whales, blue whales, Bryde’s whales, and humpback whales. Details on species sighting records and their distribution are provided in Table 1 and supplementary material 1 respectively.

Overall, ensembles created using all models performed better than individual models when compared using the AUC and CBI metrics (Figure 1). Balanced RF and MaxEnt models also performed equally well for several species. Model performance was better when species records were filtered at a 100 km scale for all species except humpback dolphins, spinner dolphins, pan-tropical spotted dolphins, Risso’s dolphins, and short-finned pilot whales, for which records were filtered at the scale of 10 km (Table 1).

**Figure 1:**
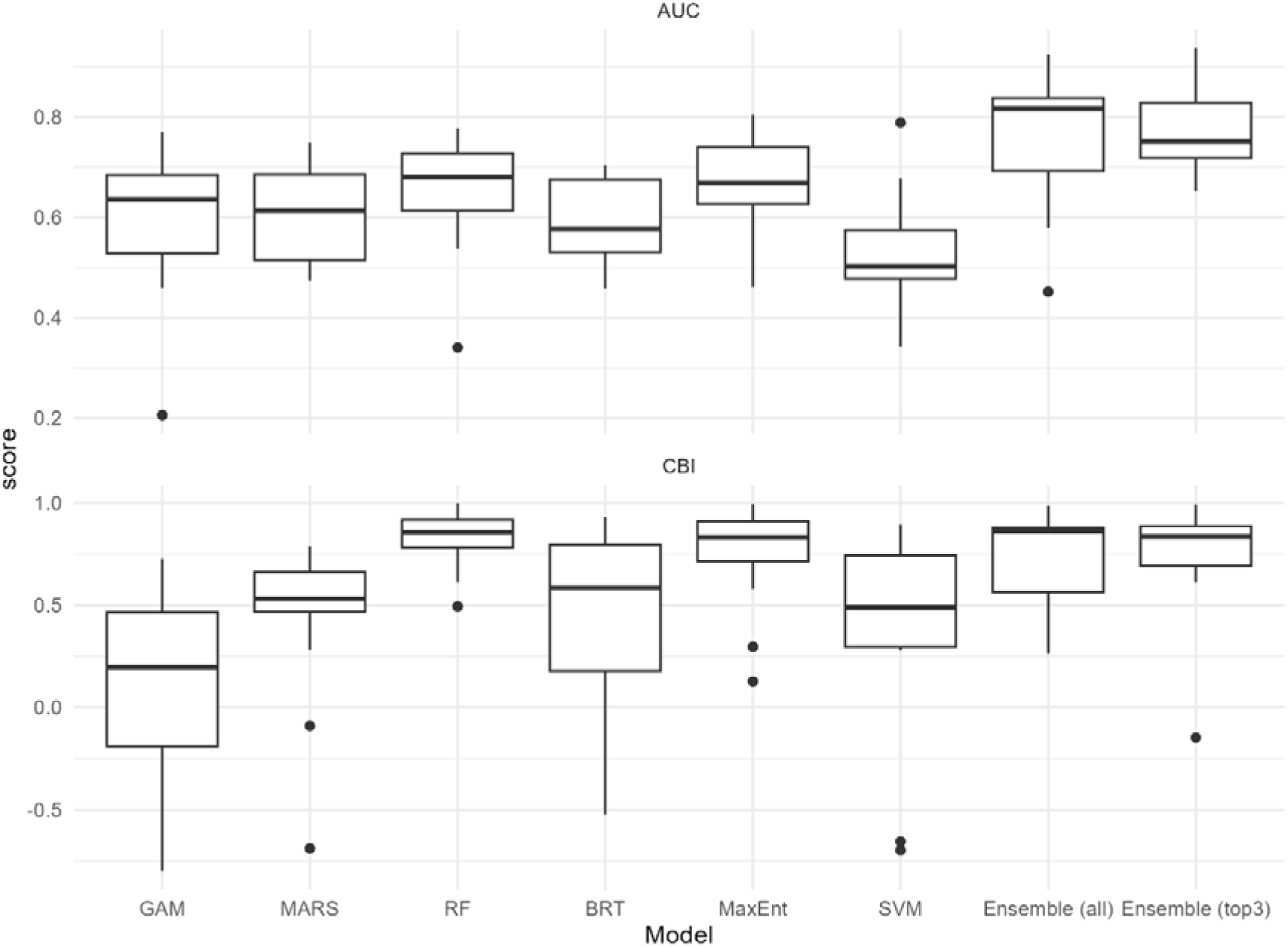
AUC and CBI score for all eight models for every species.

Generally, physiographic variables (depth, slope, and distance to land) were of greater relative importance than oceanographic variables in predicting species distributions but for some species like humpback dolphins, Irrawaddy dolphins, false killer whales, and long-beaked common dolphins, primary productivity and temperature fronts had greater percentage importance (Table A1, supplementary information). Mean SST, SST range and current velocity were highly spatially autocorrelated (>6000 km) and consequently were the least important variables.

### 3.2. Habitat suitability and occurrence

Predicted habitat suitability (Figure 2) and occurrence (supplementary material 3) varied considerably across species. Of the three nearshore species, Indo-pacific finless porpoises showed a contiguous distribution along the mainland and occurred within 20-30 km from the coastline. Their distribution overlapped greatly with humpback dolphins that had high habitat suitability along the central and southern mainland coasts. Irrawaddy dolphins occurred up to 60 km from the coast but were restricted to West Bengal and Odisha in India, and Bangladesh.

**Figure 2:**
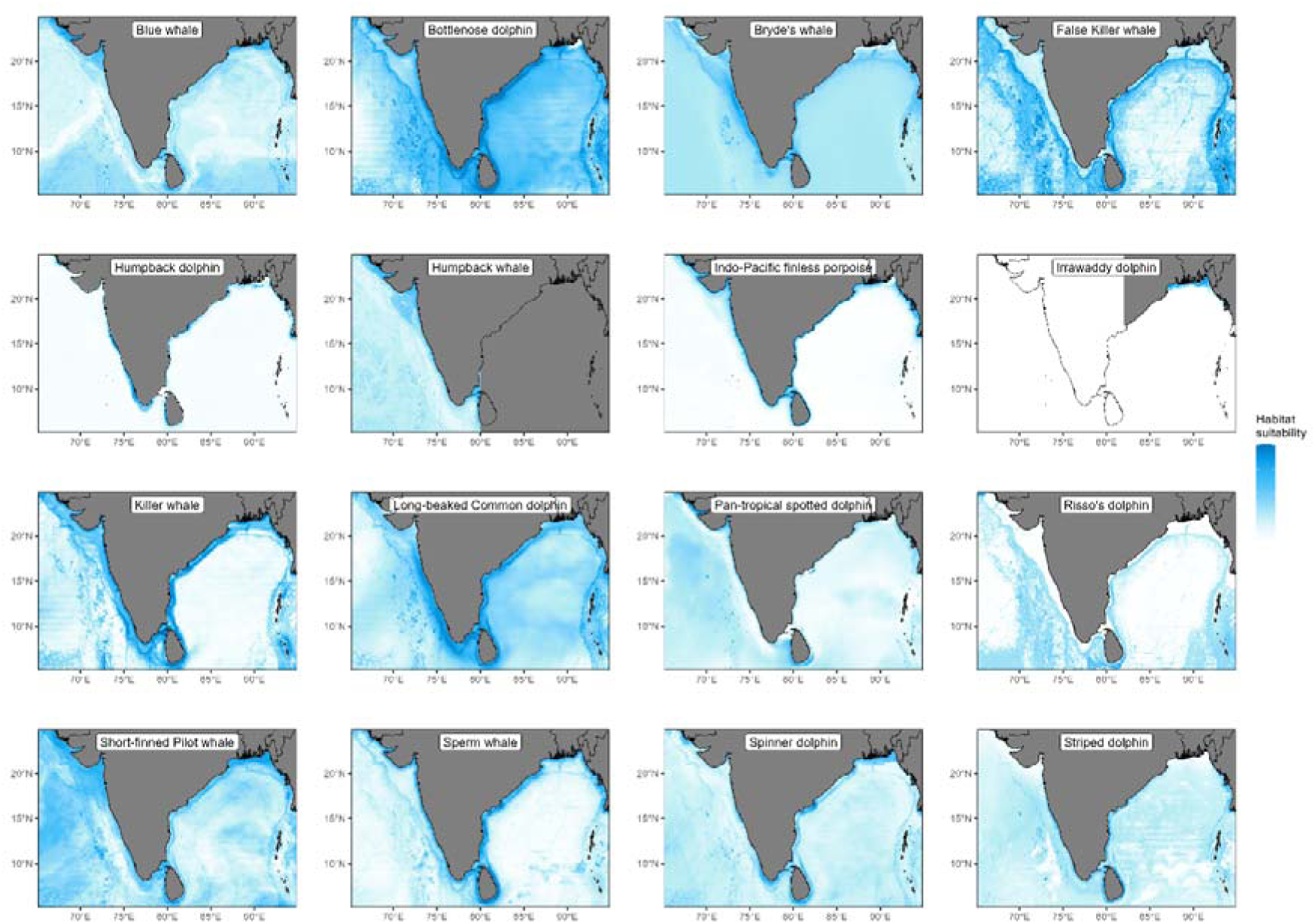
Habitat suitability maps for all 16 species groups.

The distribution of bottlenose dolphins and striped dolphins was wider and habitat suitability greater at mid/lower latitudes compared to spinner dolphins and pan-tropical spotted dolphins that occurred throughout the study area but had different patterns of habitat suitability. Long-beaked common dolphins were distributed almost throughout the study area but with few highly suitable areas offshore. On the contrary, Risso’s dolphins were distributed on and beyond the continental slope. High suitability areas for short-finned pilot whales occurred in northern Arabian sea and Bay of Bengal, and Sri Lanka. Suitable habitats for false killer whales overlapped with that of Risso’s dolphins while killer whales had suitable habitats along the Indian coast, in southern Sri Lanka and the two archipelagos of India.

Sperm whales had high habitat suitability near Sri Lanka and the east coast of India. Mid latitude regions in the Arabian sea were of high suitability for Bryde’s whales while blue whale had more suitable habitats in lower and higher latitudes. Finally, humpback whales were ubiquitous but high suitability areas occurred in the northern Arabian sea and Sri Lanka.

### 3.3 Species richness

Altogether, species richness was greatest around the continental slope and coastal regions and extended further from the coastline on the east coast compared to the west coast of India (Figure 3). The southern tip of Sri Lanka was identified as a hotspot. In offshore regions, the Arabian sea had greater species richness, specifically around the Lakshadweep archipelago, compared to the Bay of Bengal but areas around the Andaman and Nicobar islands showed high species diversity. Both these areas of high species richness were characterised by complex seabed characteristics.

**Figure 3:**
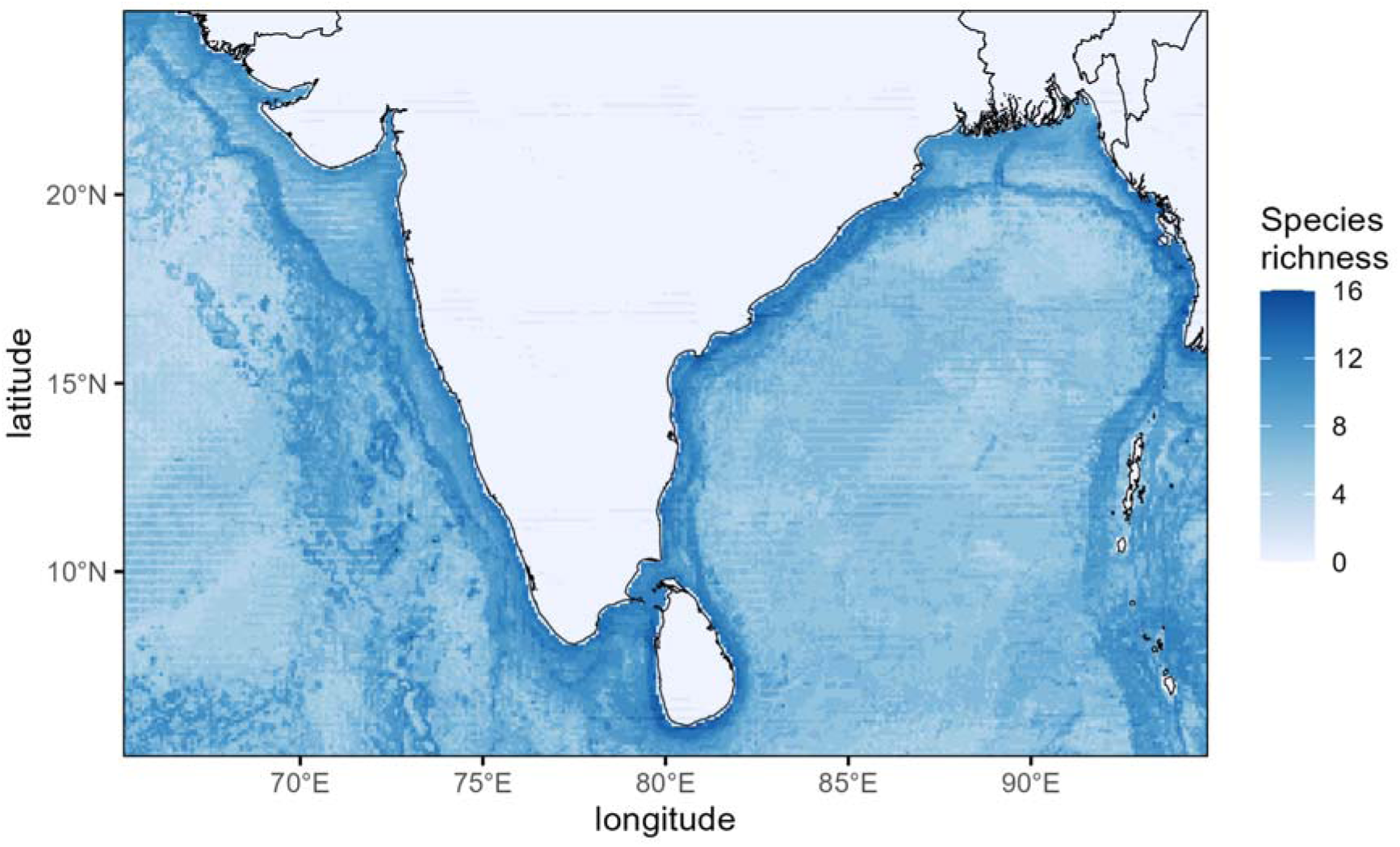
Estimated species richness in the study area.

### 3.4. Prediction validation and expert review

Areas with high uncertainty in predictions were comparable for all species, as predicted by the MESS analysis, and occurred in the Sundarbans delta of the Bangladesh region. For a few species, predictions in the offshore areas of Gujarat and Pakistan were also of relatively higher uncertainty (supplementary material 2).

We received feedback from six cetacean experts in the region. Average accuracy scores for habitat suitability predictions, as perceived by the reviewers, ranged between 4.4 and 8 (out of 10) for different species (Table 3). Predictions for nearshore species (mean accuracy score 7.5) were more accurate than that for offshore species (mean accuracy score 6.0). Generally, species with very little information available from the study area (e.g., long-beaked common dolphins, spinner dolphins) scored low. For a few species, areas of high prediction uncertainty i.e., near the Sundarbans and offshore Gujarat, were also identified as areas of unexpectedly high suitability. For eight of the 16 species groups, models ranked highest by most reviewers were also ones with the highest CBI and AUC scores. For the remaining species, reviewer-selected models differed only slightly with the models with the highest evaluation metrics (Table 3). Binary presence/absence maps for all species, highlighting over- and under-predicted areas are provided in supplementary material 3.

**Table 3:**
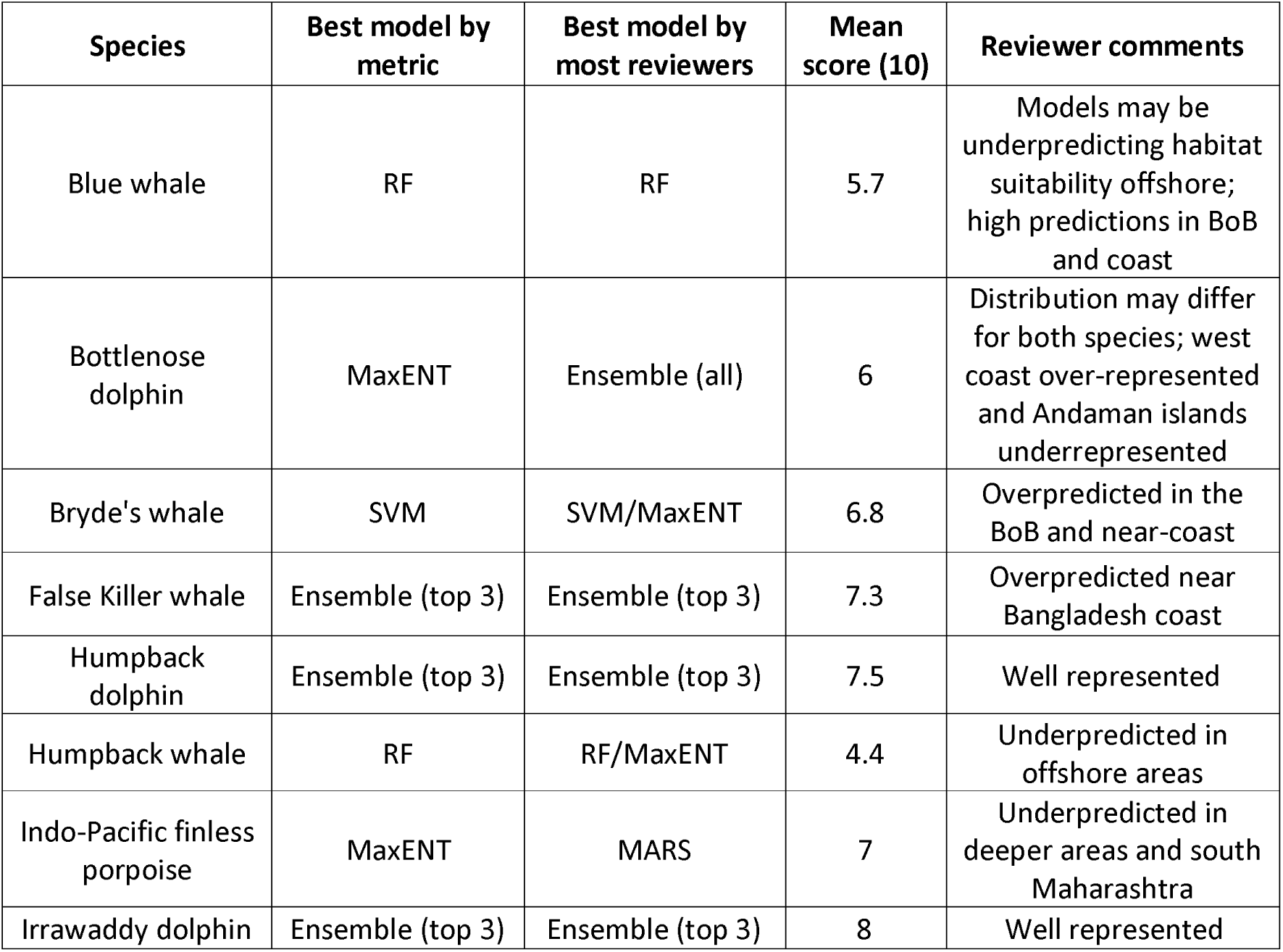
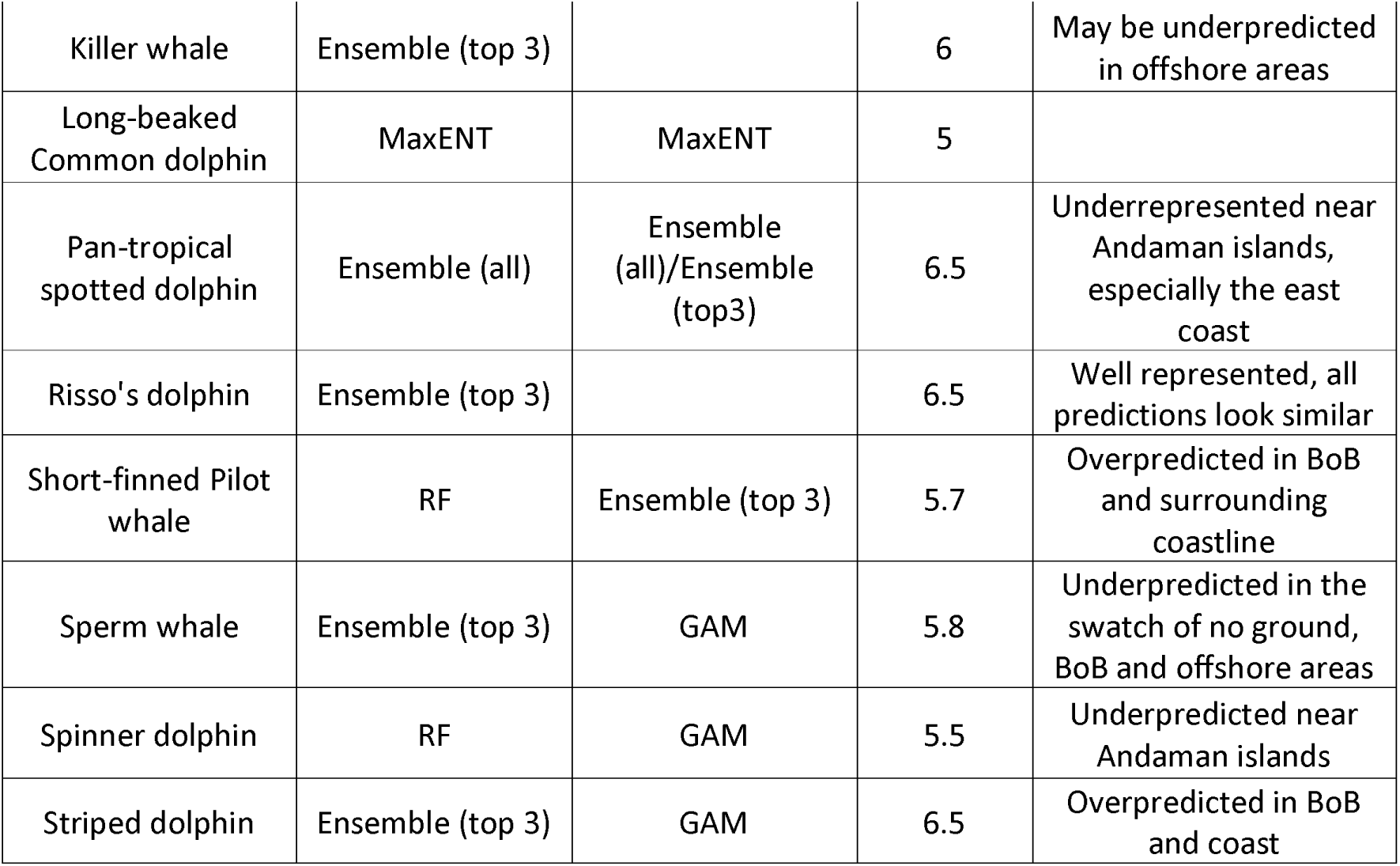
A summary of reviewer scores and responses on predicted habitat suitability and occurrence of species in the study area. All model predictions can be found in supp mat 4 and occurrence maps with over- and underpredicted areas highlighted can be found in supp mat 3. BoB refers to the Bay of Bengal.

## 4. Discussion

By combining sighting data from multiple sources, we modelled the distribution of 18 cetacean species occurring in the data-poor regions of the Arabian sea and the Bay of Bengal. Since species identity could not be resolved for the two species of humpback dolphins (*S. plumbea, S. chinensis*) and bottlenose dolphins (*T. aduncus, T. truncatus*), they were modelled as two groups, leading to 16 prediction maps. Altogether, species were predicted to occupy different regions based on their ecology and habitat requirements (Figure 2). Species richness was greatest along the mainland coasts of the Indian subcontinent and the Lakshadweep and Andaman and Nicobar archipelagos (Figure 3). The relevance of our results for further cetacean research and conservation in the region is discussed below.

### 4.1 Understanding bias and uncertainty in predictions

Based on evaluation metrics and expert review, most of our models provided reasonably accurate predictions of habitat suitability and distribution for various species. Perceived accuracy was weakly but positively correlated with sample size suggesting that for some species like blue whales (n=14 after filtering), sightings were likely missing from few areas. Sampling extent therefore had a significant effect on model performance (Rana et al., 2024), which could not be detected using evaluation metrics alone. In addition, based on the MESS index (Elith et al., 2010; supplementary material 2) regions outside the sampled covariate space i.e., with covariate values beyond the sampling range were predicted to be falsely unoccupied by several species. Altogether, our top models predicted a conservative distribution for species including blue whales, Bryde’s whales, false killer whales, short-finned pilot whales and Risso’s dolphins, especially in the Bay of Bengal. Yet, this does not refute our inference of high species richness along the east coast of India since all these species were predicted to occur in this region with high certainty, which is also confirmed by stranding reports (Dudhat et al., 2022, MMRCNI, 2024). However, species richness in the Bay of Bengal may be higher than predicted and our results advocate for increased survey effort in this region.

### 4.2 Species distribution patterns and occurrence hotspots

Depth and productivity generally defined suitable habitats for the three coastal species (Sahri et al. 2021). Of all cetaceans, humpback dolphins had a high and most evenly distributed number of records (n=157 post filtering) likely because they occur extremely close to the coast and are easily detected (Sutaria et al., 2015). Their absence in Palk Strait – the area between India and Sri Lanka was unexpected. Palk Strait has similar properties to the adjacent regions of Palk Bay and Gulf of Mannar where Indo-pacific humpback dolphins occur (Muralidharan 2019), and so, this area may be used by humpback dolphins only seasonally. The areas east and west to this pocket are occupied by Indo-pacific humpback dolphins and Indian Ocean humpback dolphins respectively and so Palk Strait, and perhaps the tip of southern India may also be a barrier restricting the movement of these species.

Within the occupied region, habitat suitability varied considerably and may reflect varying species densities. For example, humpback dolphin habitat suitability and density in the Gulf of Kutch is lower compared to the central west coast of India (Sutaria & Jefferson 2005). The west coast has more shallow and estuarine habitats compared to the east which translates to greater suitability (Parra, Schick, & Corkeron 2006).

Such differences were not identified in the distribution of Indo-pacific finless porpoises, likely due to a smaller and scattered distribution of sighting records. At fine scales, Indo-pacific finless porpoises may avoid areas with humpback dolphins (Wursig et al., 2016) and human disturbances (Fang et al., 2022). However, they are also difficult to spot during visual surveys, leading to fewer sighting records. Their distribution is likely contiguous and overlaps with humpback dolphins even in areas off Bangladesh coast where they were predicted absent (Smith et al., 2008).

Irrawaddy dolphins were predicted to occur along the coasts of Bangladesh and West Bengal, India and in the Chilika lake in Odisha. Like humpback dolphins, Irrawaddy dolphins also prefer estuaries (Peter et al., 2016) and are only present in isolated pockets of the Odisha coast (Sutaria, 2009), likely due to its bathymetry or high vessel traffic (Kreb et al., 2020).

In the case of oceanic delphinids, IUCN distribution maps suggest common bottlenose dolphins occur close to the coastline of the subcontinent while Indo-pacific bottlenose dolphins have an offshore distribution (Braulik et al., 2019; Wells et al., 2019). While this pattern is evident in our habitat suitability predictions it is not possible to segregate distributions for the two species. The distribution of bottlenose dolphins overlapped with spinner dolphins, pan-tropical spotted dolphins, and striped dolphins that showed different patterns of habitat suitability but that are comparable in body size and their ecology. Despite a high number of widely distributed sighting records, the distributions for spinner dolphins (n=171) and pan-tropical spotted dolphins (n=23) could not be predicted precisely, suggesting that delphinids may select habitat at smaller scales (Weir et al., 2012) while occupying a wider region (Oviedo et al., 2018). On the other hand, striped dolphins and long-beaked common dolphins, occurred generally at lower latitudes in both coastal and oceanic habitats. All these delphinids are likely to be sympatric with the occasional clusters of humpback dolphins and Indo-pacific finless porpoises along in the east coast, partitioning niches at finer scales or even forming mixed species groups (Gimenez et al., 2018; Syme, Kiszka & Parra, 2023).

Amongst larger, deep-water species Risso’s dolphins and false killer whales are known to prefer deeper, continental slope waters (Jefferson et al., 2014; Sahri et al., 2021), which is reflected in our predicted distribution for the species. On the continental slope, these species may overlap with short-filled pilot whales (Sankalpa et al., 2021) that occurred closer to the coast as well. Sperm whales are deep diving species and their underprediction in offshore waters highlights the need for more offshore surveys in the future.

Bryde’s whales showed a near-coastal distribution along the mainland and the Lakshadweep islands and may migrate to higher latitudes in the southern winter (June-October) (Patro et al., 2022). The distribution of blue whales was wider at lower latitudes (<10°N) and in coastal areas above 20°N in both the Arabian Sea and the Bay of Bengal. Using transfer SDMs built for the California Current system, Redfern et al. (2017) also show these areas to be of high suitability for the species, but our models fail to capture regions in the mid-latitudes (10- 20°N) which blue whales may use to migrate seasonally. Both species may use habitats in the north and south differently depending on their migratory pattern (Thums et al., 2022) and since they prefer upwelling zones, their distribution may be limited to continental shelf and slope areas (Moller et al., 2020; Ferreira et al., 2024). Their migrations may also be comparable to humpback whales that migrate from Oman towards Sri Lanka during the northern winter (December-March), while humpback whales from the southwest Indian Ocean migrate to Sri Lanka and southwest India during the southern winter (June-October) (Anderson et al., 2022). Therefore, in the Arabian Sea habitat predicted for humpback whales in the north may be occupied by the former population and in the south by the latter at different times of the year. Recent records of killer whales also suggest that they may follow a similar migratory pattern from Oman to Sri Lanka using predicted habitat, and their records from the Andaman and Nicobar islands are likely from a distinct population.

Finally, predicted patterns of species richness were broadly consistent with identified biogeographic realms in the region (Spalding et al., 2007). Specifically, richness in the east and west coasts of India differed with the former supporting larger habitats for coastal delphinids and migratory species. The narrower continental shelf in the east allowed offshore species like Risso’s dolphins to occur close to the coastline, while providing suitable habitat for several other delphinids. Cetacean diversity in the northeastern coast in the Bay of Bengal was also different from the southeastern coast with the north supporting Irrawaddy dolphins and greater habitats for humpback dolphins and Bryde’s whales. The two coasts were separated by the species-rich waters of Sri Lanka where 17 of the 18 species occurred. The Lakshadweep archipelago supported large habitats for more species compared to the west coast while excluding coastal dolphins. On the contrary, Andaman and Nicobar islands had the lowest species richness (11 of 18 species). While the two oceanic archipelagos showed marginally lower species richness compared to the mainland coast, it must be noted that of the 31 species recorded, we were only able to model the distribution of 18 species. The remaining species included members from *Ziphiidae* and *Kogiidae*, Fraser’s dolphins, rough-toothed dolphins, etc. (Table 1) which occur in deeper waters, with sparse records from the two island systems. In all regions, species occurrence and richness correlated to the presence of complex bathymetric features.

### 4.3 Future research and conservation of cetaceans in India

The paucity of sighting records for several species and our conservative distribution predictions for few others highlights the importance of focusing survey efforts on the deeper waters of the Arabian sea and the Bay of Bengal. Both these areas have high levels of shipping traffic (Hou et al., 2015) and large-scale fishing activities (Welch et al., 2022) which are primary threats to cetaceans globally, underscoring the importance of obtaining baseline information on endangered species. Closer to the coast, research activities and species records were more prevalent from the west coast of India even though the east coast showed greater species richness. While there is a need to enhance survey effort in the east coast, visual surveys must be supplemented with acoustic surveys to monitor elusive species like the Indo-pacific finless porpoise, that occurs in the same regions as humpback dolphins but had very few of sighting records after filtering (n=14). India’s Project dolphin (GoI, 2021) provides an excellent opportunity to set up baselines for species distributions to identify species rich areas and monitor future changes. Under its ambit, nation-wide surveys can be conducted annually in deep waters and perhaps seasonally along the coastline to understand occurrence as well as the coexistence mechanisms for different sympatric species. In addition, since cetaceans occur across large geographic areas, they provide excellent opportunities for fostering international cooperation through transboundary surveys (Mackelworth 2012). Finally, while mapping habitat suitability is essential, it may not always correlate with species abundance (Rullens et al., 2021). Thus, systematic surveys may be needed to detect changes in population size, providing better information on impacts of human activity on the environment (Moore 2008; Hazen et al., 2019).

There are several challenges to implementing and sustaining large-scale survey effort in developing nations like India. Alternatives like very high resolution (VHR) satellite imagery with AI tools (Kapoor et al., 2023) can be useful to cover large scale areas and understand oceanic distribution of large cetaceans. Locally, to ensure that cetacean monitoring is sustainable, it is imperative to invest in awareness and education programs that work with seafaring communities and train them in collecting opportunistic sighting records (Lodi & Tardin 2018). Several platforms to collate and organise such information like GBIF and MMRCNI are already available for public use. Individuals who spend a significant part of their time at sea including fishers, tour operators, ferry operators, etc. can be trained to collect opportunistic sighting records and upload them to open-access citizen science platforms. In addition, areas of significant interest can be monitored passively using acoustic devices to understand fine-scale cetacean habitat-use and movement. However, such large-scale programs are only bound to succeed with involvement from multiple stakeholders including government bodies.

While generating baseline information is crucial, efforts must be made parallelly to reduce impacts of known anthropogenic activities on cetaceans. Mortality from bycatch and fisheries interactions (Dudhat et al., 2022; Jog et al., 2024), especially in nearshore areas may be high and awareness programs should incorporate educating people about collecting information on stranding events. Fine-scale data and distribution models can identify hotspots of occurrence that can help prioritise areas for monitoring and management. Combined with modern and cost-effective techniques that leverage data on ocean currents to understand stranding patterns (Peltier et al., 2012), such data can also provide valuable information into bycatch patterns around the Indian coast. Similarly impacts of pollution on cetacean health can also be assessed from stranded animals while remote monitoring of offshore habitats can help identify environmental changes and understand the overlap of large whales with industrial fishing and shipping routes.

## Acknowledgements

This paper on the PhD research of IS at CES, IISc which was supported by the Prime Minister’s Research Fellowship. We thank all the reviewers including Dr. D. Panicker, Dr. G. Braulik, Ms. K. Jog, Mr. M. Sule, Ms. M. Mankeshwar, and Dr. R. Muralidharan who provided critical feedback and inputs about our models and predicted maps.

## Data availability statement

All data and codes relevant to this study will be published as open access in the GitHub repository of IS upon successful acceptance of this manuscript.

## Conflict of interest statement

The authors declare no conflict of interest.

## Notes

### Competing Interest Statement

The authors have declared no competing interest.

